# Development of genetic markers for sexing *Cannabis sativa* seedlings

**DOI:** 10.1101/2020.05.25.114355

**Authors:** Djivan Prentout, Olga Razumova, Hélène Henri, Mikhail Divashuk, Gennady Karlov, Gabriel AB Marais

**Author notes:** corresponding author: Gabriel Marais; LBBE – UMR 5558, CNRS / Université Lyon 1, campus de la doua, 69622 Villeurbanne cedex, France.

## Abstract

*Cannabis sativa* is a dioecious plant with a XY system. Only females produce cannabinoids in large amount. Efficient male removal is an important issue for the cannabis industry. We have recently identified the sex chromosomes of *C. sativa*, which opens opportunities for developing universal genetic markers for early sexing of *C. sativa* plants. Here we selected six Y-linked markers and designed PCR primers, which were tested on five hemp cultivars both dioecious and monoecious. We obtained promising results, which need to be extended using a larger number of individuals and a more diverse set of cultivars, including THC producing ones.

---

*Cannabis sativa* is a dioecious plant with XY chromosomes (Divashuk *et al.*, 2014). Only non-pollinated females produce THC and CBD (Small, 2015). Removing males as early as possible is thus a major issue for the THC/CDB industry. In *C. sativa*, however, sexual dimorphism is absent before the flowering stage as in many dioecious plants (Small, 2015; Barrett and Hough, 2013). Several methods have been developed to produce male-free crops (discussed in McKernan *et al.*, 2020). One of the most promising methods consists in identifying males before the flowering stage with Y-specific genetic markers, typically in *C. sativa* seedlings. A few genetic markers for sexing have been previously developed in *C. sativa* but accuracy or throughput need to be improved (discussed in Toth *et al.*, 2020). A recent study improved the throughput of MADC6, one of those genetic markers, using PACE - PCR Allele Competitive Extension (Törjek *et al.*, 2002; Toth *et al.*, 2020). The sex assay was conducted on 2,170 plants and 14 cultivars, and they correctly identified 98% of the females and 100% of males (in a sub-sample of 270 plants). However, MADC6 and the other available markers are located in retrotransposons and they might not be present in all *C. sativa* cultivars. Despite all these advances, we still need to develop universal genetic markers for sexing *C. sativa* seedlings.

In a recent study, we performed a genome-wide segregation analysis of *C. sativa* to identify sex-linked genes (Prentout *et al.*, 2020). We ran SEX-DETector on genotyping data of a *C. sativa* cross and identified SNPs that show sex-linkage when looking at allele transmission from parents to progeny (Muyle *et al.*, 2016; Prentout *et al.*, 2020). SEX- DETector identified more >550 sex-linked genes (Prentout *et al.*, 2020). Aligning those sex-linked genes onto a chromosome-level assembly of a *C. sativa* genome from Grassa *et al.* (2018), we found that the largest chromosome pair (number 1) was the sex chromosomes pair. Among the sex-linked genes that were identified by SEX-DETector, ∼350 were XY gene pairs and the highest synonymous divergence between X and Y sequences reached 40%. These most strongly divergent XY gene pairs are interesting because they have a very high chances to be present in all *C. sativa* populations/cultivars and thus offer a great opportunity to develop universal Y-linked genetic markers for early sexing of *C. sativa*. Here, we show the results on six such markers that have been tested on hemp cultivars.

## Methods

We used the XY gene pairs with the highest synonymous divergence identified in Prentout *et al.* (2020), for which CDS length was greater than 300 pb. We aligned the sequences with CLC workbench tool ‘create alignment’ (version 8.0.1, see https://www.quiagenbioinformatics.com) and uses the tool ‘Design Primer’ for primer design. For each gene pair, we aligned the Y-linked sequence from our cross (male × female of hemp cultivar Zenitsa, see Prentout *et al.*, 2020) with three X-linked sequences, one of which was from our cross and the other two from a Purple Kush cultivar (THC producer) female and from a Finola cultivar (hemp) female (Van bakel *et al.*, 2011). This increased the probability of detecting X/Y fixed differences (shared by all individuals of the species) instead of X/Y polymorphism (specific to some populations). To design the primers, we selected coding regions with high X-Y divergence. We kept those with at least two fixed differences between X-linked and Y-linked sequences, and avoided complementary forward and reverse primers to avoid association between our primers during PCR.

We used the *in silico* PCR tool from the Van bakel lab (http://genome.ccbr.utoronto.ca/) to test the size of the amplicons, and verify that a unique region of the genome was amplified. Our primers were tested *in silico* on both the Purple Kush and the Finola genomes. As both genomes are female genomes, if the X-linked primers resulted in amplification but the Y-linked did not, we validated the primers as sufficiently divergent between the X-linked and Y-linked copies.

We then tested our markers *in vitro* by PCR. Plant material was obtained from different cultivars and DNA was extracted. DNA isolation was performed from young leaves as described by Doyle and Doyle (1990) with some modifications. The extracting buffer contained 100 mM Tris-HCl (pH = 8.0), 20 mM EDTA (pH = 8.0), 2 M NaCl, 1.5% CTAB, 1.5% PVP and 0.2% β-mercaptoethanol. A 15 mM ammonium acetate solution in 75% ethanol was used for DNA washing. The PCR program for the all primers contained the following steps: 95 °C for 5 min followed by 35 cycles of 94 °C for 30 sec, 55 °C for 1 min, and 72 °C for 1 min and a final step of 72°C for 7 min. The markers were tested on three dioecious hemp cultivars (Zenitsa, Viktoria and Ekaterinodar) and two monoecious hemp cultivars (1147|16 and Maria). As the monoecious plants are XX (Razumova *et al.*, 2014) the Y-linked markers primers are not supposed to amplify. They can thus serve as control.

## Results

Using the approach described in Methods, we selected thirteen XY gene pairs and twelve autosomal genes. We designed primer pairs for both X-linked and Y-linked sequences (XY gene pairs) for the autosomal sequence (autosomal genes). All the thirteen genetic Y-linked and twelve autosomal markers were validated *in silico* (see Methods). The primers were first tested *in vitro* on Zenitsa plant material (also used in Prentout *et al.*, 2020). From this preliminary test, we selected six Y-linked and one autosomal markers that amplified well with the primers that we designed and the PCR conditions that we set.

Table 1 shows the results of PCR experiments on those Y-linked and one autosomal markers in five different cultivars (three dioecious and two monoecious). Except for the Ekaterinodar cultivar, the Y-linked primers amplified in 100 % of males and 0 % of females and monoecious cultivars. The autosomal control markers amplified in all tested individuals. Amplicon size ranged from 174 to 305 bp.

**Table 1:** PCR validation of six Y-linked markers. Results are given for six Y-linked primer pairs and 1 autosomal (control) primer pair. Five different cultivars have been tested: three dioecious (Zenitsa, Viktoria, Ekaterinodar) and two monoecious (‘1447|16’ and Maria). Sample sizes are indicated with cultivars’ names. We had roughly 50:50 males/females in all cultivars. The percent of individuals that amplified is given for each marker in each cultivar. For monoecious cutlivars, plants are considered females because their genotype is XX. Therefore, a PCR in monoecious male is not applicable (na).

| Cultivars | Y-linked marker 8 |  | Y-linked marker 10 |  | Y-linked marker 21 |  | Y-linked marker 23 |  | Y-linked marker 36 |  | Y-linked marker 37 |  | Autosomal marker 16 |  |
| --- | --- | --- | --- | --- | --- | --- | --- | --- | --- | --- | --- | --- | --- | --- |
|  | M | F | M | F | M | F | M | F | M | F | M | F | M | F |
| **Zenitsa**<br>(n = 26) | 100 | 0 | 100 | 0 | 100 | 0 | 100 | 0 | 100 | 0 | 100 | 0 | 100 | 100 |
| **Viktoria**<br>(n = 8) | 100 | 0 | 100 | 0 | 100 | 0 | 100 | 0 | 100 | 0 | 100 | 0 | 100 | 100 |
| **Ekaterinodar**<br>(n = 14) | 80 | 11 | 100 | 0 | 80 | 11 | 0 | 0 | 80 | 11 | 80 | 0 | 100 | 100 |
| **1447\|16 (XX)**<br>(n = 2) | na | 0 | na | 0 | na | 0 | na | 0 | na | 0 | na | 0 | na | 100 |
| **Maria (XX)**<br>(n = 6) | na | 0 | na | 0 | na | 0 | na | 0 | na | 0 | na | 0 | na | 100 |

## Discussion

We identified six very promising markers, which amplified only in males for the dioecious cultivars (except for Ekaterinodar) and did not amplify in monoecious cultivars. Results in Ekaterinodar were more ambiguous. We observed some amplification in one female for three markers: 8, 21 and 36 (see Table 1). Moreover, the markers 8, 21, 36 and 37 amplified only in 80% of the males for this cultivar. Marker 23 did not amplify in Ekaterinodar. Only marker 10 had amplification in 100% males and 0% females. It is unclear why results were not as good in this cultivar compared to others. It should be noted however that combining the results of the six markers identified correctly male and female plants in Ekaterinodar.

More generally, sample sizes were small and the rates of success are thus only very rough. More tests are thus necessary. We need to test our markers on a much larger number of plants, from a much more diversified set of cultivars (e.g. including THC producers).

## References

Barrett SC, Hough J. 2013. Sexual dimorphism in flowering plants. J Exp Bot 64(1): 67–82.

Divashuk, M. G., Alexandrov, O. S., Razumova, O. V., Kirov, I. V., & Karlov, G. I. (2014). Molecular cytogenetic characterization of the dioecious Cannabis sativa with an XY chromosome sex determination system. PloS one, 9(1).

Doyle, J.J., Doyle, J.L. (1990). Isolation of plant DNA from fresh tissue. Focus 12:13–15

Grassa, C. J., Wenger, J. P., Dabney, C., Poplawski, S. G., Motley, S. T., Michael, T. P., … & Weiblen, G. D. (2018). A complete Cannabis chromosome assembly and adaptive admixture for elevated cannabidiol (CBD) content. BioRxiv, 458083.

McKernan, K. J., Helbert, Y., Kane, L. T., Ebling, H., Zhang, L., Liu, B., … & Concepcion, G. (2020). Sequence and annotation of 42 cannabis genomes reveals extensive copy number variation in cannabinoid synthesis and pathogen resistance genes. BioRxiv.

Muyle, A., Käfer, J., Zemp, N., Mousset, S., Picard, F., & Marais, G. A. (2016). SEX-DETector: a probabilistic approach to study sex chromosomes in non-model organisms. Genome biology and evolution, 8(8), 2530–2543.

Prentout, D., Razumova, O., Rhoné, B., Badouin, H., Henri, H., Feng, C., … & Marais, G. A. (2020). An efficient RNA-seq-based segregation analysis identifies the sex chromosomes of Cannabis sativa. Genome Research, 30(2), 164–172.

Razumova, O. V., Alexandrov, O. S., Divashuk, M. G., Sukhorada, T. I., & Karlov, G. I. (2016). Molecular cytogenetic analysis of monoecious hemp (Cannabis sativa L.) cultivars reveals its karyotype variations and sex chromosomes constitution. Protoplasma, 253(3), 895–901.

Small, E. (2015). Evolution and classification of Cannabis sativa (marijuana, hemp) in relation to human utilization. The botanical review, 81(3), 189–294.

Törjék, O., Bucherna, N., Kiss, E., Homoki, H., Finta-Korpelová, Z., Bócsa, I., … & Heszky, L. E. (2002). Novel male-specific molecular markers (MADC5, MADC6) in hemp. Euphytica, 127(2), 209–218.

Toth, J. A., Stack, G. M., Cala, A. R., Carlson, C. H., Wilk, R. L., Crawford, J. L., … & Smart, L. B. Development and validation of genetic markers for sex and cannabinoid chemotype in Cannabis sativa L. GCB Bioenergy.

Van Bakel, H., Stout, J. M., Cote, A. G., Tallon, C. M., Sharpe, A. G., Hughes, T. R., & Page, J. E. (2011). The draft genome and transcriptome of Cannabis sativa. Genome biology, 12(10), R102.

